# *Rhizobium desertarenae* sp. nov., isolated from the Saline Desert Soil from the Rann of Kachchh, India

**DOI:** 10.1101/2020.07.03.186106

**Authors:** Mitesh Khairnar, Ashwini Hagir, Avinash Narayan, Kunal Jain, Datta Madamwar, Yogesh Shouche, Praveen Rahi

**Author notes:** Corresponding author details: Praveen Rahi, National Centre for Microbial Resource, National Centre for Cell Science, Pune, Maharashtra 411007, India, Email ID. The GenBank accession numbers for the 16S rRNA gene sequences of strain ADMK78^T^ is MK942856. The genome sequence has been deposited in GenBank under the accession number CP058350-CP058352. The type strain is available with different culture collections under the accession numbers MCC 3400^T^; KACC 21383^T^; and JCM 33657^T^.

## Abstract

A novel bacterial strain designated ADMK78^T^ was isolated from the saline desert soil. The cells were rod-shaped, Gram-negative, and non-motile. The strain ADMK78^T^ grows best at 28°C and pH 7.0 and can tolerate up to 2% (w/v) NaCl. Based on 16S rRNA gene phylogeny, the strain ADMK78^T^ belongs to the genus *Rhizobium*, with the highest similarity to *Rhizobium wuzhouense* W44^T^ (98.7%) and *Rhizobium ipomoeae* shin9-1^T^ (97.9%). Core-genes based phylogenetic analysis revealed that the strain ADMK78^T^ forms a distinct branch in between *Rhizobium ipomoeae* shin9-1^T^ and *Rhizobium selenitireducens* BAA-1503^T^. The average nucleotide identity of ADMK78^T^ was less than 82%, to members of the family *Rhizobiaceae*. The genomic DNA G+C content of strain ADMK78^T^ is 58.6 mol%. The major fatty acids of strain ADMK78^T^ were C_18:0_ and C_18:1 ω7c_. The strain ADMK78^T^ showed differences in physiological, phenotypic, and protein profiles estimated by MALDI-TOF MS to its closest relatives. Based on the phenotypic, chemotaxonomic properties, and phylogenetic analyses, the strain ADMK78^T^ could be distinguished from the recognized species of the genus *Rhizobium*. It is suggested to represent a novel species of this genus, for which the name *Rhizobium desertarenae* sp. nov. is proposed. The type strain is ADMK78^T^ (=MCC 3400^T^; KACC 21383^T^; JCM 33657^T^).

## Introduction

The genus *Rhizobium* is a large group of bacteria, and most species are known for their symbiotic fixation of nitrogen within the root nodules of leguminous plants. The family *Rhizobiaceae* has now been classified into 16 genera [1,2]. *Rhizobium* is one of the main genera in the family *Rhizobiaceae*, and it was first proposed in 1889 [3,4]. Presently, the genus *Rhizobium* comprises 90 recognized species (https://lpsn.dsmz.de/genus/rhizobium). Members of the genus *Rhizobium* are characterized as Gram-stain-negative, non-spore-forming, rod-shaped, aerobic, chemo-organotrophic, and possess C_18:1_ and C_18:1 ω7c_ as the predominant fatty acid and have a DNA G+C content of between 55 and 66 mol% [5,6]. Non-symbiotic and free-living members of *Rhizobium* have been found in various soils, including the rhizosphere [7,8], bioreactor [9], and beach sand [10].

### Isolation and Ecology

During the investigations on bacterial diversity of the saline desert soil collected (23.7337° N, 69.8597° E) from the Rann of Kachchh, India, a bacterial strain ADMK78^T^, was isolated on Zobell Marine Agar. Rann of Kachchh is reputed to be the largest salt desert in the world and is a transitional area between marine and terrestrial ecosystems [11]. The region experiences diagonal fluctuations of average temperatures with a high temperature of 50 °C during summers and drops below freezing during winters. Due to the hot and hypersaline environment, there is a vast possibility of identifying novel microbes with high economic and industrial potential. The newly isolated strain ADMK78^T^ was maintained on Zobell Marine Agar at 37 °C, and preserved at -80 °C as a suspension in 20% (v/v) glycerol and by lyophilization with 20% (w/v) skimmed milk. The pure culture was processed for MALDI-TOF MS-based identification, and a comparison of the MALDI-TOF MS spectrum of ADMK78^T^ with the Biotyper 3.0 database resulted in no reliable identification.

### 16S RNA phylogeny

High-quality genomic DNA was extracted from the strain following the JGI protocol version 3 for bacterial genomic DNA isolation using CTAB [12]. The 16S rRNA gene sequence was amplified using universal primers (27f: 5’-AGAGTTTGATCCTGGCTCAG-3’ and 1492r: 5’-TACGGCTACCTTGTTACGACTT-3’) according to the methods described by Gulati *et al*. [13], and the amplified product was directly sequenced using the ABI PRISM Big Dye Terminator v3.1 Cycle Sequencing kit on a 3730xl Genetic Analyzer (Applied BioSystems, Thermo Scientific, USA). The similarity search for the 16S rRNA gene sequence of strain ADMK78^T^ was performed against the type strains of prokaryotic species in the EzBioCloud’s valid species database [14]. The strain ADMK78^T^ showed the highest similarity to *Rhizobium wuzhouense* W44^T^ (98.7%) and followed by *Rhizobium ipomoeae* shin9-1^T^ (97.9%). The 16S rRNA gene sequence of strain ADMK78^T^ was used as queries to closely related gene sequences using the NCBI BLASTn tool [15] and the non-redundant nucleotide database. Higher than 99% sequence similarity was recorded for two sequences. The first one was of Alpha proteobacterium (EU770254.1) associated with *Microcystis aeruginosa* culture, and the second one was of *Ciceribacter* sp. strain AIY3W (MH463946.2) isolated from low salinity lakes on Tibetan Plateau. Multiple alignments of sequences of strain ADMK78^T^ and its nearest neighbours retrieved from EzBioCloud’s server and NCBI GenBank, and phylogenetic analyses were performed using MEGA software (version 7.0) [16]. Bootstrap values were determined based on 1000 replications. The 16S rRNA gene sequence of *Bradyrhizobium japonicum* USDA 6^T^ was used as an outgroup. The strain ADMK78^T^ formed a separate branch along with the Alpha proteobacterium and *Ciceribacter* sp. AIY3W (Fig. 1). The valid species of the genus *Ciceribacter* were placed in a distinct group, which was far from the group formed by ADMK78^T^, Alpha proteobacterium, and *Ciceribacter* sp. AIY3W. The overall topologies were similar for the phylogenetic trees obtained with the ML, MP and NJ methods. *Rhizobium ipomoeae* shin9-1^T^ appears to be the closest phylogenetic neighbour to the group formed by ADMK78^T^, Alpha proteobacterium, and *Ciceribacter* sp. AIY3W in all three trees constructed by different methods.

**Fig. 1:**
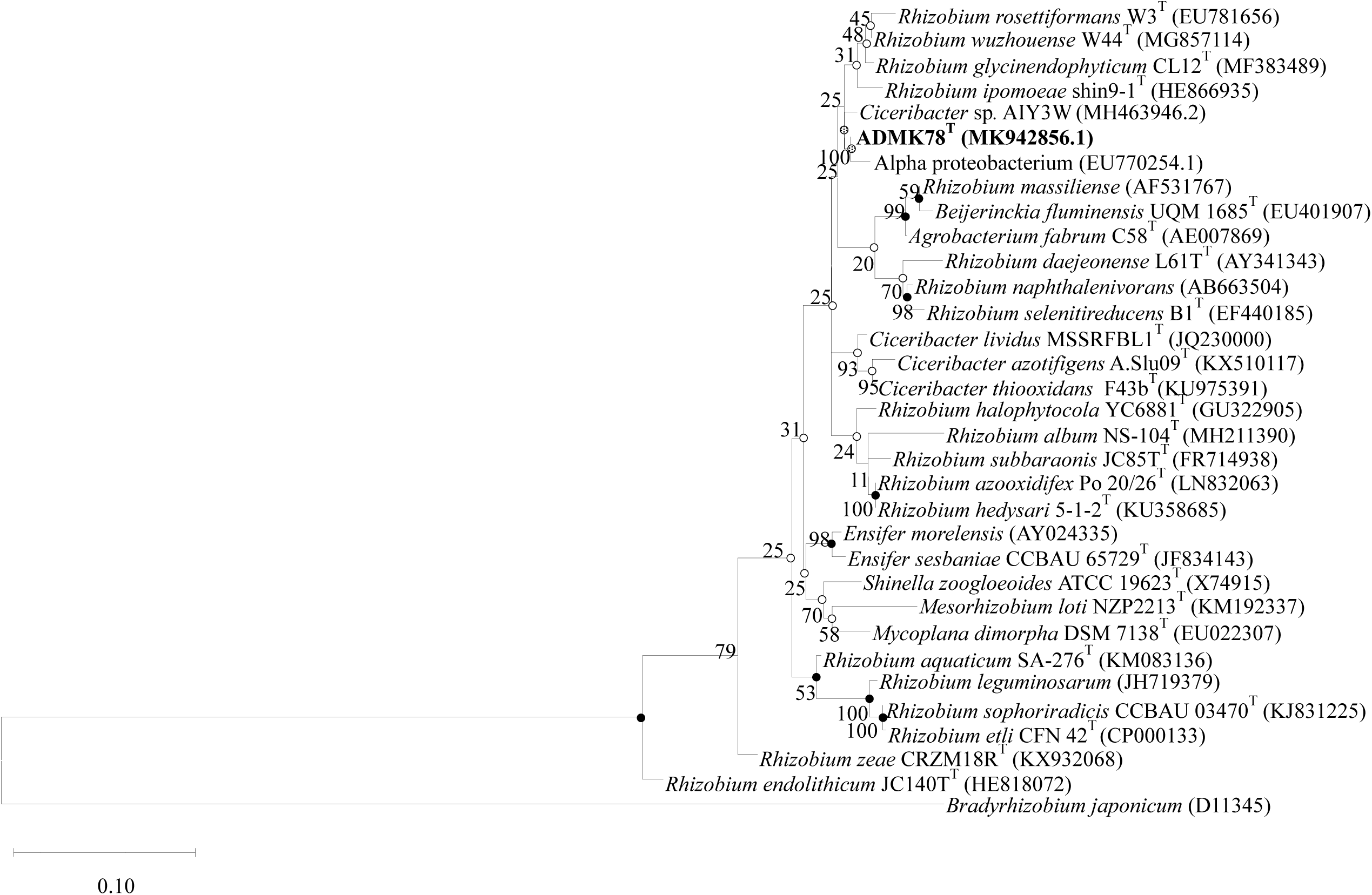
Phylogenetic tree was inferred by using the Maximum Likelihood method and Tamura-Nei model. The percentage of trees in which the associated taxa clustered together is shown next to the branches. Initial tree(s) for the heuristic search were obtained automatically by applying Neighbor-Join and BioNJ algorithms to a matrix of pairwise distances estimated using the Tamura-Nei model, and then selecting the topology with superior log likelihood value. Empyt circles indicate branches of the tree that were also recovered using the neighbour-joining method, circles with dots indicate recovery with the maximum-parsimony method, and black filled circles indicate that all three methods recovered the corresponding nodes. *Bradyrhizobium japonicum* USDA 6^T^ (D11345) was used as an outgroup. Bar, 0.1 substitutions per nucleotide position.

### Genome Features

Genome sequencing was performed using a hybrid approach of two platforms, first on an Illumina MiSeq platform with 2 × 250 bp v2 chemistry, followed by sequencing with Oxford Nanopore Technology (ONT) on a minION platform. The Nanopore reads were assembled using Canu v. 2.0 [17] with default settings. The overlaps between the ends of circular contigs were identified using NUCmer v. 3.1 [18] and removed using a custom Perl script. Two rounds of polishing was performed using the paired-end Illumina reads. In each round the Illumina reads were mapped to the genome assembly using bowtie2 v. 2.3.4.1 [19] with default parameters, followed by polishing using Pilon v. 1.23 [20] with default settings. The genome sequence quality of strain ADMK78^T^ was as per the genome standards proposed by Chun et al. [21], and the detailed genome features are provided in Table 1. The genome of ADMK78^T^ had a size of 4,342,374 bp, which is smaller than the genome size of symbiotic members of *Rhizobium* and in the range of the sizes of the non-symbiotic strains of *Rhizobium* (Table 1). It consisted of a circular chromosome of 3,590,542 bp and two circular plasmids of 708,533 and 43,299 bp. The overall genome sequencing coverage for the strain ADMK78^T^ was 147.5x, with an N50 value of 3,590,542 bp. Whole-genome sequences were annotated using the RAST [22] web server (http://rast.nmpdr.org/rast.cgi). The genome of strain ADMK78^T^ contains 4377 protein-coding sequences (CDS), of which 64 genes assigned to the stress response functions, like heat and cold shock, hyperosmotic stress, and protection from reactive oxygen. We could not find any nitrogen-fixation and nodulation genes in the genome of strain ADMK78^T^.

**Table 1:**
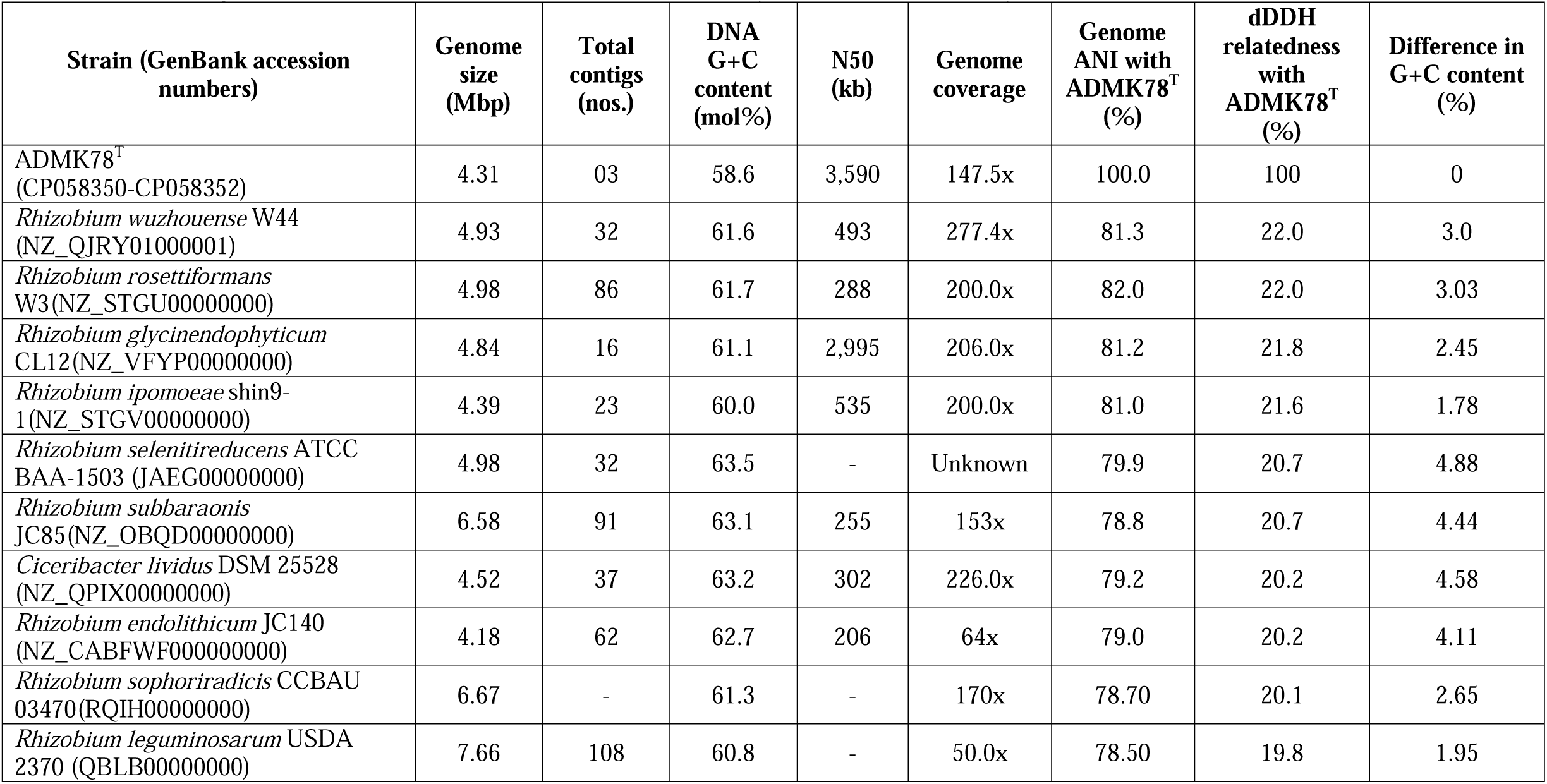
Standard genome features of strain ADMK78^T^ and related type strains of the family *Rhizobiaceae*

The bacterial core gene-based phylogenetic analysis was carried out using the UBCG pipeline [23] from the concatenated sequences of 92 core genes extracted by UBCG, and a maximum-likelihood phylogenetic tree was inferred using RAxML version 8.2.8 [24] with the GTRGAMMA model and 100 bootstrap replications. The Average Nucleotide Identity (ANI) was determined between strain ADMK78^T^ and closely related strains of the *Rhizobiaceae* family using FastANI [25], and a heatmap representation of the calculated ANI values was constructed using DisplayR (https://www.displayr.com/). The genome-based phylogenetic analysis placed strain ADMK78^T^ as an independent branch, with *Rhizobium ipomoeae* shin9-1^T^ as the closest neighbour and followed by *Rhizobium rosettiformans* W3^T^, *Rhizobium wuzhouense* W44^T^ and *Rhizobium glycinendophyticum* CL12^T^ (Fig. 2). The highest ANI values of ADMK78^T^ was for *Rhizobium rosettiformans* W3 (82%), followed by *Rhizobium wuzhouense* W44^T^ (81.3%) and *Rhizobium ipomoeae* shin9-1^T^ (81%) (Fig. S1; Table 1). The dDDH relatedness values of ADMK78^T^ with the reference strains were below 22% (Table 1), suggesting the strain ADMK78^T^ is a novel species [19]. The genomic DNA G+C content of strain ADMK78^T^ was 58.6 mol%, which is well within the range (i.e., 55-66 mol %) of the genus *Rhizobium* [4].

**Fig. 2:**
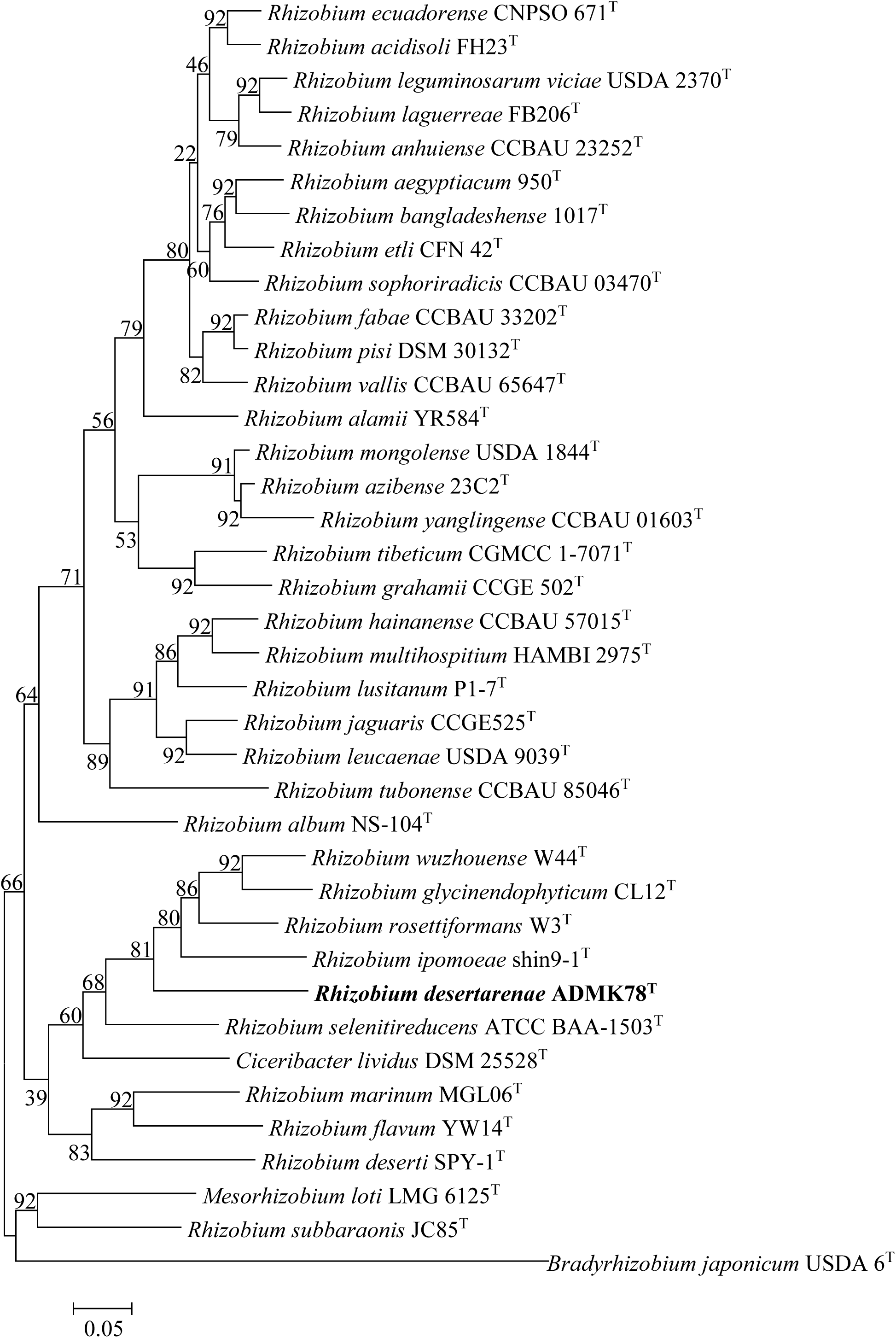
Phylogenetic tree inferred by the UBCG phylogenomics pipeline using the concatenated alignment of 92 core genes, of strain ADMK78^T^ and its closely related taxa. Percentage of bootstrap values are given at branching points. Bar, 0.05 substitution per position.

### Physiology and Chemotaxonomy

The type strains of *Rhizobium ipomoeae* shin9-1^T^ and *Ciceribacter lividus* MSSRFBL1^T^ were obtained from the Deutsche Sammlung von Mikroorganismen und Zellkulturen (DSMZ) and the BCCM/LMG Bacteria Collection, Belgium (LMG), respectively. Both type strains were used as reference strains and evaluated together under identical experimental conditions to those for strain ADMK78^T^.

For analysis of chemotaxonomic features, the strain was grown on Zobell Marine Agar at 28 °C, and cell biomass was harvested after 24 h. Preparation and analysis of fatty acid methyl esters were performed as described by Sasser [25] using the Microbial Identification System (MIDI) and the Microbial Identification software package (Sherlock version 6.1; MIDI database, TSBA6). The primary fatty acids detected in strain ADMK78^T^ were C_18:1 ω7c_ and C_18:0_ (Table S1). Small proportions of C_16:0,_ C_18:1 ω7c 11-methyl,_ C_18:0 3OH,_ C_20:1 ω7c,_ and summed feature 2 (C_12:0 aldehyde_) were also detected for strain ADMK78^T^. The strain ADMK78^T^ exhibit higher proportions of C_18:0,_ similar to *Ciceribacter lividus* MSSRFBL1^T^ (Table S1), which was missing in *Rhizobium wuzhouense* W44^T^ [26] and present in relatively lower amounts in *Rhizobium ipomoeae* shin9-1^T^. A relatively higher proportion of C_18:1 ω7c 11-methyl_ and C_20:1 ω7c_in the fatty acids profile of strain ADMK78^T^, differentiate it from the reference strains (Table S1).

Whole-cell proteins were extracted using ethanol/formic acid after 24 h growth on TSA, to generate the Mean Spectral Profile (MSP). The proteins ranging from 2-20 KDa were analyzed by matrix-assisted laser desorption/ionization time-of-flight mass spectrometer (MALDI-TOF MS) autoflex speed (Bruker Daltonik GmbH, Germany) [27]. A total of 27 replicate spectra were used to generate a new MSP of strain [28], which was compared with MSPs of the reference strains *Rhizobium ipomoeae* shin9-1^T^ and *Ciceribacter lividus* MSSRFBL1^T^ generated during this study following same procedure. The mean spectra profile of strain ADMK78^T^ has 33 unique peaks in comparison to the closely related taxa out of a total of 70 peaks (Table S2), which attributed to the discrimination of ADMK78^T^ from the closely related species.

Morphological, physiological, and biochemical tests for strain ADMK78^T^ were performed on Zobell Marine Agar plates incubated under aerobic conditions. The colonies of strain ADMK78^T^, were circular and translucent on Zobell Marine Agar (Fig. S3). Gram-staining (K001 and K004, Himedia, India) was used following the manufacturer’s instructions. Hanging drop technique was used to check the motility. Scanning electron microscopy was performed to observe cell morphology as described in Rahi *et al*. [29]. These analyses revealed that strain ADMK78^T^ is a rod-shaped bacterium with cell size ranging from 0.3-0.5×1.5-2 µm, Gram stain negative, (Fig. S2) and non-motile. Oxidase disc (DD018, Himedia, India) was used for testing oxidase activity, and catalase activity was determined by bubble formation in a 3□% (v/v) H_2_O_2_ solution. The strain was positive for both oxidase and catalase. Growth at different temperatures (4, 10, 15, 20, 28, 37, 45 and 55 °C), NaCl concentrations [0-2% (w/v) at 0.5% intervals] and pH values (4.0-11.0 at 1.0 pH unit intervals) was examined after incubation in Zobell Marine broth for 7 days in automated microbial growth analyzer (Bioscreen C, OY Growth Curves, Finland). The initial pH of the inoculation broth was adjusted using 1 M HCl and 1 M NaOH. The strain ADMK78^T^ grows at 10-45 °C (optimum 28°C), pH ranging from 4-10 (optimum 7.0), and NaCl concentration tolerance up to 2%.

Biochemical characteristics, enzyme activities, and oxidation/or reduction of carbon sources were performed using the API 20E and API ZYM systems (07584D and 25200, bioMérieux, France) and Biolog GN III system (OmniLog, Biolog, USA) following manufacturers’ instructions. The Biolog test showed that out of the substrates present in the GENIII BIOLOG microplate, ADMK78^T^ showed activity for 71 substrates, of which a weak reaction was recorded for four substrates (Table S3). The strain ADMK78^T^ can be distinguished from its closest phylogenetic neighbours based on features listed in Table 2.

**Table 2.** Differentiating characteristics of strain ADMK78 ^T^ in comparison to its closest phylogenetic neighbours. strain 1, *Rhizobium* sp. ADMK78 ^T^; 2, *Rhizobium wuzhouense* W44^T^; 3,*Rhizobium ipomoeae* sp. shin9-1^T^; 4, *Ciceribacter lividus* MSSRFBL1^T^

| Characteristics | 1 | 2* | 3 | 4 |
| --- | --- | --- | --- | --- |
| Isolation source | Saline soil (Desert) | Roots of *Oryza officinalis* | Field | Rhizosphere soil of chickpea |
| Colony colour | Cream | Cream | Cream | Bluish black |
| pH range for growth | 4.0-11(7) | 5-8 | 7.0-9.0(7) | 6.0-8.5(7) |
| Temperature range for growth (°C) (optimum) | 10-45 (28) | 15-40 | 10-45 (30) | 10-45 (28) |
| NaCl range for growth (%) (optimum) | 0-2.0 (1.5) | 0-2.0 | 0-3.0 (1.5) | 0.5-1.5 (1) |
| D-raffinose | - | + | - | + |
| rifamycin sv | - | - | + | + |
| Gelatin | - | + | - | - |
| L-arginine | + | - | + | + |
| β-galactosidase | - | + | + | - |
| glucosidase | - | ND | + | - |
| DNA G+C content (mol%) | 58.6 | 61.6 | 60.0 | 63.2 |

\*Data from Tao Yuan et al., [27]

The genotypic and phenotypic data generated for the strain ADMK78^T^ revealed that the strain represents a novel species in the genus *Rhizobium*, for which the name *Rhizobium desertarenae* sp. nov. is proposed.

### Description of *Rhizobium desertarenae* sp. nov

***Rhizobium desertarenae*** (de.sert.a.re’nae. L. neut. n. *desertum* desert; L. fem. n. *arena* sand; N.L. gen. n. *desertarenae* of desert sand).

Cells are Gram-negative, straight rods with round ends (0.3-0.5×1.5-2 µm), and non-motile. Colonies grown on Zobell Marine Agar are 1-3 mm in diameter, circular, raised with an entire margin, and translucent opacity. The optimal temperature for growth is 28 °C, and the optimal pH is 7.0. Growth occurs in the absence of NaCl with up to 2% tolerance in Zobell Marine broth. It is oxidase and catalase positive. The strain showed positive results in Biolog GN III analyses for utilization of D-maltose, D-trehalose, D-cellobiose, D-gentiobiose, sucrose, D-turanose, α-D-lactose, D-melibiose, β-methyl-d-glucoside, D-salicin, N-acetyl-D-glucosamine, N-acetyl-β-D-mannosamine, N-acetyl-D-galactosamine, α-D-glucose, D-mannose, D-fructose, D-galactose, D-fucose, L-fucose, L-rhamnose, inosine, D-sorbitol, D-mannitol, D-arabitol, myo-inositol, glycerol, D-glucose-6-phosphate, D-fructose-6-phosphate, D-aspartic acid, Glycyl-L-proline, glycyl-L-proline, L-alanine, L-arginine, L-aspartic acid, L-glutamic acid, L-histidine, L-pyroglutamic acid, L-serine, pectin, D-galacturonic acid, D-gluconic acid, D-glucuronic acid, glucuronamide, mucic acid, D-saccharic acid, p-hydroxy-phenylacetic acid, D-lactic acid methyl easter, L-lactic acid, citric acid, α-keto-glutaric acid, D,L-malic acid, bromo-succinic acid, Tween 40, γ-amino-butryric acid, α-hydroxy-butyric acid, α-hydroxy-D,L butyric acid, α-keto-butyric acid, acetoacetic acid, propionic acid, acetic acid, formic acid, sodium lactate, tetrazolium violet and blue, nalidixic acid, lithium chloride (Table S3). Positive results in API ZYM strips for leucine arylamidase, trypsin, naphthol-AS-BI-phosphohydrolase, α-glucosidase, N-acetyl-ß-glucosaminidase activities (Table S4). C_18:0_ and C_18:1 ω7c_ are the predominant cellular fatty acids. The DNA G+C content of the type strain is 58.6 mol%.

The type strain ADMK78^T^ (=MCC 3400^T^; KACC 21383^T^; JCM 33657^T^) was isolated from saline desert sand collected from the Kutch District of Gujarat, India. The GenBank sequence accession number of the genome sequence is CP058350-CP058352, and 16S rRNA gene sequence of strain ADMK78^T^ is MK942856.

## Acknowledgments

The authors thank Prof. Aharon Oren, Department of Plant and Environmental Sciences, The Hebrew University of Jerusalem, Israel and Prof. Bernhard Schink, Department of Biology, University of Konstanz, Germany, for their expert suggestions concerning the correct species name, species epithet and Latin etymology.

## Ethical Statement

The experiments reported in this manuscript did not involve human participants and/or animals.

## ABBREVIATIONS

CDS: coding sequence
CTAB: Cetyl trimethylammonium bromide;
MALDI-TOF MS: Matrix-Assisted Laser Desorption/Ionization Time-of-Flight Mass Spectrometer;
MSP: Mean Spectral Profile;
UBCG: Up-to-date bacterial core genes.

