## Supplementary data for "*Rhizobium desertarenae* sp. nov., isolated from the Saline Desert Soil from the Rann of Kachchh, India"

**Corresponding author details**:

Praveen Rahi,

**Table S1:** A comparative account of fatty acid composition (%) of strain 1, *Rhizobium* sp. ADMK78 ^T^; 2, *Rhizobium wuzhouense* W44^T^; 3,*Rhizobium ipomoeae* shin9-1^T^; 4, *Ciceribacter lividus* MSSRFBL1^T^

| **Fatty Acids** | **1** | **2*** | **3** | **4** |
| --- | --- | --- | --- | --- |
| C_11 : 0 10-methyl_ | ND | 0.1 | ND | ND |
| C_13 : 0 12-methyl_ | ND | 0.1 | ND | ND |
| C_15:0 iso_ | 0.2 | ND | ND | ND |
| C_15:1_ _ω8c_ | ND | ND | 0.2 | ND |
| **C_16:0_** | **1.1** | **2.5** | **1.0** | **3.03** |
| C_16:0_ _3OH_ | ND | 0.1 | 0.3 | 0.23 |
| C_17:1_ _ω7c_ | 0.9 | ND | 0.7 | ND |
| C_17:0_ | 0.4 | 0.2 | ND | 0.31 |
| C_18:0 iso_ | 0.5 | ND | 0.9 | ND |
| **C_18:1_ _ω9c_** | **0.56** | **ND** | **ND** | **ND** |
| C_18 : 1 ω12c_ | ND | 0.1 | ND | ND |
| C_18:1_ _ω5c_ | 0.3 | ND | 0.3 | 0.31 |
| **C_18:0_** | **10.2** | **ND** | **2.3** | **9.7** |
| **C_18:1_ _ω7c 11-methyl_** | **2.4** | **ND** | **0.3** | **0.8** |
| C_18 : 1 ω6c 11-methyl_ | ND | 2.1 | ND | ND |
| **C_18:0_ _3OH_** | **1.0** | **2.3** | **1.07** | **2.57** |
| **C_19:0_ _cyclo ω8c_** | **ND** | **ND** | **ND** | **17.88** |
| C_19 : 1 ω9c_ | ND | 0.1 | ND | ND |
| **C_20:1_ _ω7c_** | **2.3** | **ND** | **0.55** | **0.80** |
| C_20:2_ _ω6,9c_ | ND | 2.0 | ND | 0.52 |
| C_20:0_ | 0.2 | ND | ND | ND |
| Summed feature 1 | ND | 0.3 | ND | ND |
| **Summed feature 2 (C_12:0 aldehyde_)** | **3.7** | **0.5** | **4.5** | **4.5** |
| Summed feature 3 (C_16:1 ω7c/16:1 ω6c_) | 0.3 | ND | 0.9 | 0.8 |
| Summed feature 7 (C_19:1 ω7c/19:1 ω6c_) | 0.3 | ND | ND | 0.5 |
| **Summed feature 8 (C_18:1 ω7c_)** | **75.9** | **89.0** | **87.3** | **58.4** |

ND, Not detected; *data from Tao Yuan et al., 2018 [27]

**Table S2**: List of unique protein biomarker peaks (m/z) of strain ADMK78^T^ in comparison to the closely related taxa.

| **S.No.** | ****Rhizobium* sp.**  **ADMK78 ^T^** | ***Rhizobium ipomoeae***  **shin9-1^T^** | ***Ciceribacter lividus***  **MSSRFBL1^T^** |
| --- | --- | --- | --- |
| 1 | 3083 | 3083 | 3088 |
| 2 | 3404 | 3411 | 3405 |
| 3 | 3606 | 3609 | ­­ |
| 4 | **3737** | ­­ | ­­ |
| 5 | **4411** | ­­ | ­­ |
| 6 | **4499** | ­­ | 4501 |
| 7 | 4597 | ­­ | ­­ |
| 8 | 4618 | 4626 | 4623 |
| 9 | 4668 | 4664 | 4666 |
| 10 | **4698** | ­­ | ­­ |
| 11 | **4799** | ­­ | ­­ |
| 12 | **4901** | ­­ | ­­ |
| 13 | 4921 | ­­ | 4930 |
| 14 | **4938** | ­­ | ­­ |
| 15 | 4964 | 4964 | 4965 |
| 16 | 4975 | 4967 | ­­ |
| 17 | 4997 | 4979 | 4987 |
| 18 | 5122 | ­­ | 5115 |
| 19 | 5151 | 5134 | 5141 |
| 20 | 5170 | ­­ | 5177 |
| 21 | **5199** | ­­ | ­­ |
| 22 | **5212** | ­­ | ­­ |
| 23 | **5238** | ­­ | ­­ |
| 24 | 5614 | ­­ | 5619 |
| 25 | 5641 | 5634 | ­­ |
| 26 | **5661** | ­­ | ­­ |
| 27 | 5681 | 5674 | 5681 |
| 28 | 6270 | 6274 | ­­ |
| 29 | 6582 | 6588 | ­­ |
| 30 | 6638 | 6627 | ­­ |
| 31 | 6658 | ­­ | 6657 |
| 32 | **6771** | ­­ | ­ |
| 33 | 6791 | ­­ | 6792 |
| 34 | 6805 | ­­ | 6807 |
| 35 | 6831 | 6823 | ­­ |
| 36 | 7143 | ­­ | 7147 |
| 37 | 7165 | 7174 | ­­ |
| 38 | 7197 | 7196 | ­­ |
| 39 | 7211 | 7216 | ­­ |
| 40 | 7238 | 7244 | ­­ |
| 41 | **7341** | ­­ | ­­ |
| 42 | 7365 | 7373 | ­­ |
| 43 | **7409** | ­­ | ­­ |
| 44 | **7436** | ­­ | ­­ |
| 45 | **7457** | ­­ | ­­ |
| 46 | 7472 | ­­ | 7468 |
| 47 | 7494 | 7489 | 7488 |
| 48 | 7549 | 7546 | ­­ |
| 49 | **7569** | ­­ | ­­ |
| 50 | **7682** | ­­ | ­­ |
| 51 | 7712 | ­­ | 7719 |
| 52 | 8209 | 8211 | 8215 |
| 53 | 8897 | 8898 | ­­ |
| 54 | **8935** | ­­ | ­­ |
| 55 | **8959** | ­­ | ­­ |
| 56 | **8993** | ­­ | ­­ |
| 57 | **9027** | ­­ | ­­ |
| 58 | **9152** | ­­ | ­­ |
| 59 | **9189** | ­­ | ­­ |
| 60 | **9231** | ­­ | ­­ |
| 61 | **9333** | ­­ | ­­ |
| 62 | 9390 | ­­ | 9383 |
| 63 | 9414 | ­­ | 9414 |
| 64 | **9952** | ­­ | ­­ |
| 65 | **9990** | ­­ | ­­ |
| 66 | **10294** | ­­ | ­­ |
| 67 | **11232** | ­­ | ­­ |
| 68 | **11273** | ­­ | ­­ |
| 69 | **11313** | ­­ | ­­ |
| 70 | **13301** | ­­ | ­­ |

*values from MSP list generated in this study; similar peak values (±10 Da) for reference strains obtained from the MSP lists in the biotyper database. Protein biomarker peaks highlighted in bold are specific to strain ADMK78^T^.

**Table S3:** Utilization pattern of different substrates by strain ADMK78 ^T^, *Rhizobium ipomoeae* shin9-1^T^, and *Ciceribacter lividus* MSSRFBL1^T^ on Biolog GN III plates, and *Rhizobium wuzhouense* W44^T^ (API)( Tao Yuan et al., 2018)

| **Biolog Substrate** | **1** | **2** | **3** | **4*** |
| --- | --- | --- | --- | --- |
| Dextrin | 0 | 0 | 1 | ND |
| D-maltose | 1 | 1 | 1 | 1 |
| D-trehalose | 1 | 1 | 1 | 1 |
| D-cellobiose | 1 | 1 | 1 | 1 |
| D-gentiobiose | 1 | 1 | 1 | 1 |
| Sucrose | 1 | 1 | 1 | 1 |
| D-turanose | 1 | 1 | 1 | 1 |
| Stachyose | 0 | 0 | 1 | ND |
| Positive control | 1 | 1 | 0.5 | ND |
| pH 6 | 1 | 1 | 0 | ND |
| pH 5 | 0 | 0 | 0 | ND |
| D-raffinose | 0 | 0 | 1 | 1 |
| α-d-lactose | 1 | 1 | 1 | 1 |
| D-melibiose | 1 | 0.5 | 1 | 1 |
| b-methyl-d-glucoside | 1 | 1 | 1 | ND |
| D-salicin | 1 | 1 | 1 | 1 |
| N-acetyl-d-glucosamine | 1 | 1 | 1 | ND |
| N- acetyl-b-d-mannosamine | 1 | 1 | 1 | ND |
| N-acetyl-d-galactosamine | 1 | 1 | 1 | ND |
| N-acetyl neuraminic acid | 0 | 1 | 0 | ND |
| NaCl 1% | 1 | 1 | 0.5 | ND |
| NaCl 4% | 1 | 0 | 0 | ND |
| NaCl 8% | 0 | 0 | 0 | ND |
| α-D-glucose | 1 | 1 | 1 | 1 |
| D-mannose | 1 | 1 | 1 | 1 |
| D-fructose | 1 | 1 | 1 | 1 |
| D-galactose | 1 | 1 | 1 | 1 |
| 3-methyl glucose | 0 | 0 | 0 | ND |
| D-fucose | 1 | 1 | 1 | 1 |
| L-fucose | 1 | 1 | 1 | 1 |
| L-rhamnose | 1 | 1 | 1 | 1 |
| Inosine | 1 | 1 | 1 | ND |
| Sodium lactate 1% | 1 | 1 | 1 | ND |
| Fusidic acid | 0 | 0 | 0 | ND |
| D-serine | 0 | 0 | 0 | ND |
| D-sorbitol | 1 | 1 | 1 | 1 |
| D-mannitol | 1 | 1 | 1 | 1 |
| D-arabitol | 1 | 1 | 1 | 1 |
| Myo-inositol | 1 | 1 | 1 | 1 |
| Glycerol | 1 | 1 | 1 | 1 |
| D-glucose-6-PO4 | 1 | 0.5 | 1 | ND |
| D-fructose-6-PO4 | 1 | 1 | 1 | ND |
| D-aspartic acid | 0.5 | 0 | 0 | ND |
| D-serine | 0 | 0 | 0 | ND |
| Troleandomycin | 0 | 0 | 0 | ND |
| Rifamycin sv | 0 | 1 | 1 | ND |
| Minocycline | 0 | 0 | 0 | ND |
| Gelatin | 0 | 0 | 0 | 1 |
| Glycyl-L-proline | 1 | 1 | 1 | ND |
| L-alanine | 1 | 1 | 1 | ND |
| L-arginine | 1 | 1 | 1 | 1 |
| L-aspartic acid | 1 | 1 | 1 | ND |
| L-glutamic acid | 1 | 1 | 1 | ND |
| L-histidine | 1 | 0 | 0 | ND |
| L-pyroglutamic acid | 0.5 | 1 | 1 | ND |
| L-serine | 1 | 1 | 1 | ND |
| Lincomycin | 0 | 1 | 1 | ND |
| Guanidine HCl | 0 | 0 | 1 | ND |
| Niaproof 4 | 0 | 0 | 0 | ND |
| Pectin | 1 | 1 | 1 | ND |
| D-galacturonic acid | 1 | 1 | 1 | ND |
| L-galactonic acid lactone | 0 | 1 | 1 | ND |
| D-gluconic acid | 0.5 | 1 | 1 | ND |
| D-glucuronic acid | 1 | 1 | 1 | ND |
| Glucuronamide | 1 | 1 | 1 | ND |
| Mucic acid | 1 | 1 | 0 | ND |
| Quinic acid | 0 | 1 | 1 | ND |
| D-saccharic acid | 1 | 1 | 0 | ND |
| Vancomycin | 0 | 0 | 0 | ND |
| Tetrazolium violet | 0.5 | 0.5 | 0 | ND |
| Tetrazolium blue | 1 | 1 | 1 | ND |
| p-hydroxy-phenylacetic acid | 1 | 1 | 0 | ND |
| Methyl pyruvate | 0 | 1 | 1 | ND |
| D-lactic acid methyl easter | 1 | 1 | 0 | ND |
| L-lactic acid | 1 | 1 | 1 | ND |
| Citric acid | 1 | 1 | 1 | ND |
| α-keto-glutaric acid | 1 | 1 | 0 | ND |
| D-malic acid | 1 | 1 | 0 | ND |
| L-malic acid | 1 | 1 | 0 | ND |
| Bromo-succinic acid | 1 | 1 | 0 | ND |
| Nalidixic acid | 1 | 1 | 1 | ND |
| Lithium chloride | 1 | 0.5 | 1 | ND |
| Potassium tellurite | 0 | 1 | 1 | ND |
| Tween 40 | 1 | 0 | 0 | ND |
| G-amino-butryric acid | 1 | 1 | 1 | ND |
| α-hydroxy-butyric acid | 1 | 1 | 1 | ND |
| α-hydroxy-D,L Butyric Acid | 1 | 1 | 1 | ND |
| α-keto-butyric acid | 1 | 1 | 1 | ND |
| Acetoacetic acid | 1 | 0 | 1 | ND |
| Propionic acid | 1 | 1 | 1 | ND |
| Acetic acid | 1 | 1 | 1 | ND |
| Formic acid | 1 | 1 | 1 | ND |
| Aztreonam | 0 | 0 | 1 | ND |
| Sodium butyrate | 0 | 0 | 1 | ND |
| Sodium bromate | 0 | 0 | 0 | ND |

**Strains**: 1, ADMK78 ^T^; 2, *Rhizobium ipomoeae* shin9-1^T^; 3, *Ciceribacter lividus* MSSRFBL1^T^; 4, *Rhizobium wuzhouense* W44^T^ *data from Tao Yuan et al., 2018 [27]

*1, positive; 0, negative;0.5, weekly positive.

**Table S4**: Enzyme activity of strain ADMK78 ^T^ using API-Zym test strip.

| **Enzyme assayed for** | **1** | **2** | **3** |
| --- | --- | --- | --- |
| Alkaline phosphatase | **-** | **+** | **+** |
| Esterase | - | - | - |
| Esterase lipase | - | - | - |
| Lipase | - | - | - |
| Leucine acrylamidase | **+** | **+** | **+** |
| Valine acrylamidase | - | - | - |
| Cysteine acrylamidase | - | - | - |
| Trypsin | + | + | + |
| α-chymotrypsin | - | - |  |
| Acid phosphatase | **-** | **-** | **+** |
| Naphthol AS-BI-phosphohydrolyase | **+** | **-** | **+** |
| α-galactosidase | - | + | - |
| ß-galactosidase | **-** | **-** | **-** |
| ß-glucuronidase | - | - | - |
| α-glucosidase | + | + | + |
| glucosidase | - | + | - |
| N-acetyl-ß-glucosaminidase | + | + | + |
| α-mannosidase | - | - | - |
| α-fucosidase | - | - | - |

Strains: 1, ADMK78 ^T^; 2, *Rhizobium ipomoeae* shin9-1^T^; 3, *Ciceribacter lividus* MSSRFBL1^T^;

+, positive; -, negative.

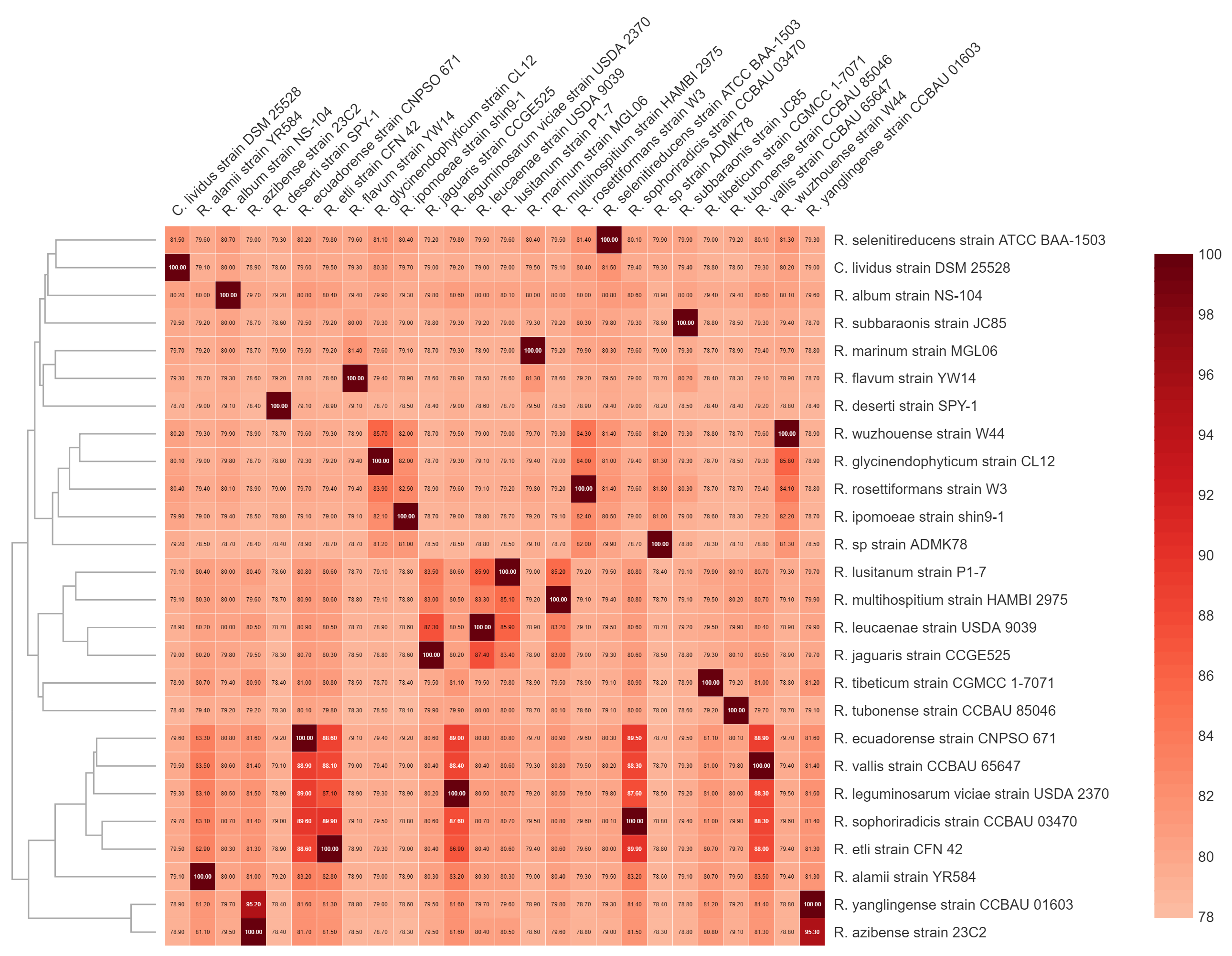

**Fig. S1**: Heatmap based on the ANI values of strain ADMK78^T^ and related type strains of

the family *Rhizobiaceae*

b Gram staining

a Gram staining

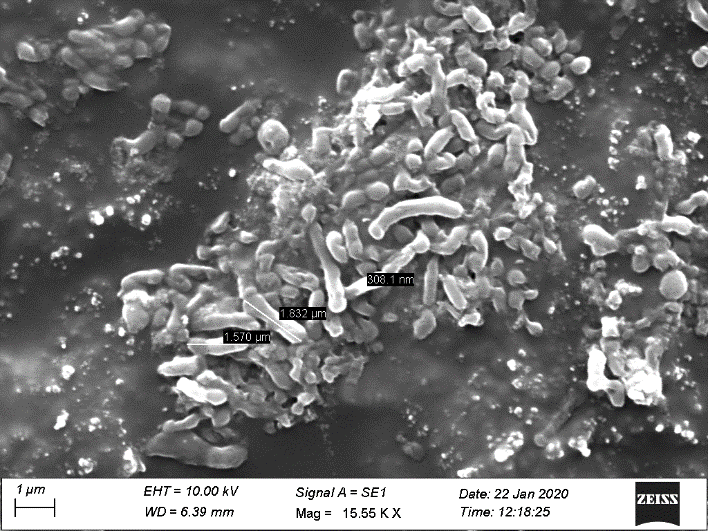

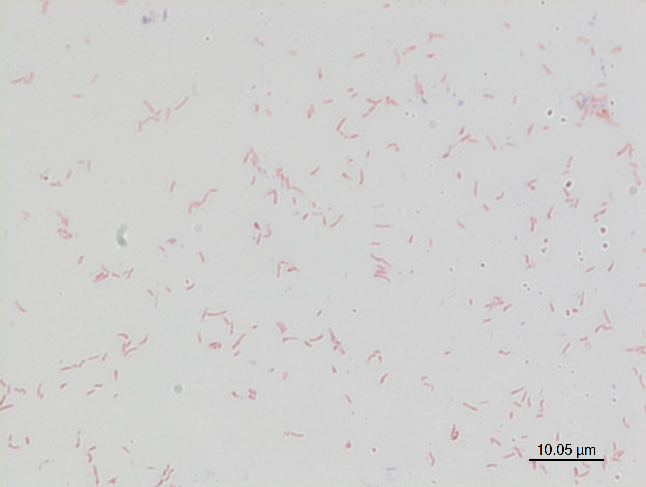

**Fig. S2**. Photomicrographs of strain ADMK78 ^T^ (a)scanning electron microscopy and (b) Gram staining

**Gram Character**: Gram Negative

**Motility**: Non-motile

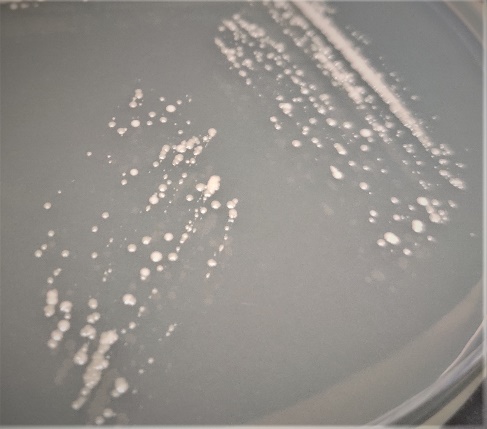

**Fig. S3**. Colonies of strain ADMK78^T^ after 24 h of incubation at 28 °C as observed on Zobell Marine Agar
